# Accounting for fragments of unexpected origin improves transcript quantification in RNA-seq simulations focused on increased realism

**DOI:** 10.1101/2021.01.17.426996

**Authors:** Avi Srivastava, Mohsen Zakeri, Hirak Sarkar, Charlotte Soneson, Carl Kingsford, Rob Patro

**Affiliations:** New York Genome Center and NYU Center for Genomics and Systems Biology, New York City, NY, USA; Department of Computer Science and Center for Bioinformatics and Computational Biology, University of Maryland, College Park, MD, USA; Harvard Medical School, Boston, Massachusetts, USA; Friedrich Miescher Institute for Biomedical Research, Basel, Switzerland; SIB Swiss Institute of Bioinformatics, Basel, Switzerland; Computational Biology Department, School of Computer Science, Carnegie Mellon University, Pittsburgh, PA, USA

**Keywords:** RNA-seq, alignment, mapping, quantification, gene expression

## Abstract

Transcript and gene quantification is the first step in many RNA-seq analyses. While many factors and properties of experimental RNA-seq data likely contribute to differences in accuracy between various approaches to quantification, it has been demonstrated (1) that quantification accuracy generally benefits from considering, during alignment, potential genomic origins for sequenced fragments that reside outside of the annotated transcriptome.

Recently, Varabyou et al. (2) demonstrated that the presence of transcriptional noise leads to systematic errors in the ability of tools — particularly annotation-based ones — to accurately estimate transcript expression. Here, we confirm the findings of Varabyou et al. (2) using the simulation framework they have provided. Using the same data, we also examine the methodology of Srivastava et al.(1) as implemented in recent versions of salmon (3), and show that it substantially enhances the accuracy of annotation-based transcript quantification in these data.

## Introduction

In their recently published paper, Varabyou et al. (2) introduce a new simulation procedure for RNA-seq data that aims to improve the realism of simulations by including fragments arising from alternative transcript start or end sites observed in experimentally acquired tissue, as well as by including simulated fragments arising from different types of “transcriptional noise” present in experimental samples. The paper both introduces the simulation framework, and assesses the performance of StringTie2 (4), paired with HISAT2 (5), as well as the annotation-based quantification methods salmon (3) and kallisto (6). They demonstrate that, as one would expect, accounting for tissue-specific variation in transcripts as well as transcriptional noise — fragments arising from transcribed RNA that does not “match” annotated isoforms — can lead to improved quantification of transcript abundance. Yet, in tissues and organisms for which a well-annotated transcriptome exists, and in which we expect most biologically meaningful expression variation to occur among annotated transcripts, the use of annotation-based quantification tools (3, 6) is likely to be an appealing option.

In the analysis performed by Varabyou et al. (2), these annotation-based methods are mapping the fragments only to the transcriptome, and are therefore unable to account for fragments that are incompatible with the annotated transcript models or that arise from other loci in the genome. Even fragments arising from annotated regions may arise in unexpected ways (*e.g*. a known UTR attached to a transcript for which it was not previously annotated, or shifts in annotated transcriptional start or termination sites). This has also been shown by Soneson et al. (7) to present a challenge for annotation-based transcript quantification tools, where the authors also introduced a measure to help identify expression estimates that may be impacted by this effect. In many cases, these issues can manifest as the false positive expression of annotated transcripts, as discussed in (1). We hypothesize that the alignment methodology recommended in Srivastava et al. (1) would help to ameliorate some of the effects of sequenced fragments arising in a manner not consistent with, or fully explained by, the annotated transcriptome. Unfortunately, the manuscript of Varabyou et al. (2) was submitted before the paper by Srivastava et al. (1) was published, so that the improved methodology explored in the latter paper was not considered in the former publication.

Here, we reassess the performance of annotation-based methods considered in Varabyou et al. (2), and evaluate the methods under some other common metrics not reported in that paper, while also including the improved methodology introduced by Srivastava et al. (1). In that work, the authors introduce two main improvements to salmon (3); first, they introduce an improved alignment method called selective-alignment and second, they provide a mechanism for including the target genome into the salmon index as a “decoy” sequence that can be aligned against to help avoid otherwise spurious mappings to annotated transcripts. We find that these additions improve the quantification performance of the annotation-based approach, salmon (3), in the simulations evaluated herein, leading to a considerably reduced number of false positive transcripts discovered, improved F_1_ scores, improved Spearman correlations with ground truth abundances, and reduced mean absolute relative differences compared to quantification when considering alignment and mapping only to the annotated transcript sequence.

## Results

For all 6 configurations (see Methods) of the quantification tools, we computed the distribution of Spearman correlation coefficients with the true fragment counts used in the simulations (Fig. 1), the distribution of the F_1_ scores of the quantification results with the truly expressed transcripts (Fig. 2), the distribution of the false positive and false negative counts (Fig. 3) and the mean absolute relative differences (MARDs) (Fig. 4) across all 30 “real” samples and all 30 “all” samples. The “real” and “all” labels follow the nomenclature of Varabyou et al. (2) and both are simulated from StringTie2 (4)-assembled transcripts. The “real” samples include only simulated fragments from transcripts compatible (as determined via their intron chain) with annotated transcripts, while the “all” samples include fragments simulated from splicing, intronic and intergenic transcriptional noise. Thus, it is worth noting that one may expect the “real” samples to be somewhat cleaner than experimental data and the “all” samples to include types of transcriptional noise that are likely to occur in many experimental samples. For more details about the samples and the definitions of the metrics, see Methods below.

**Fig. 1.**
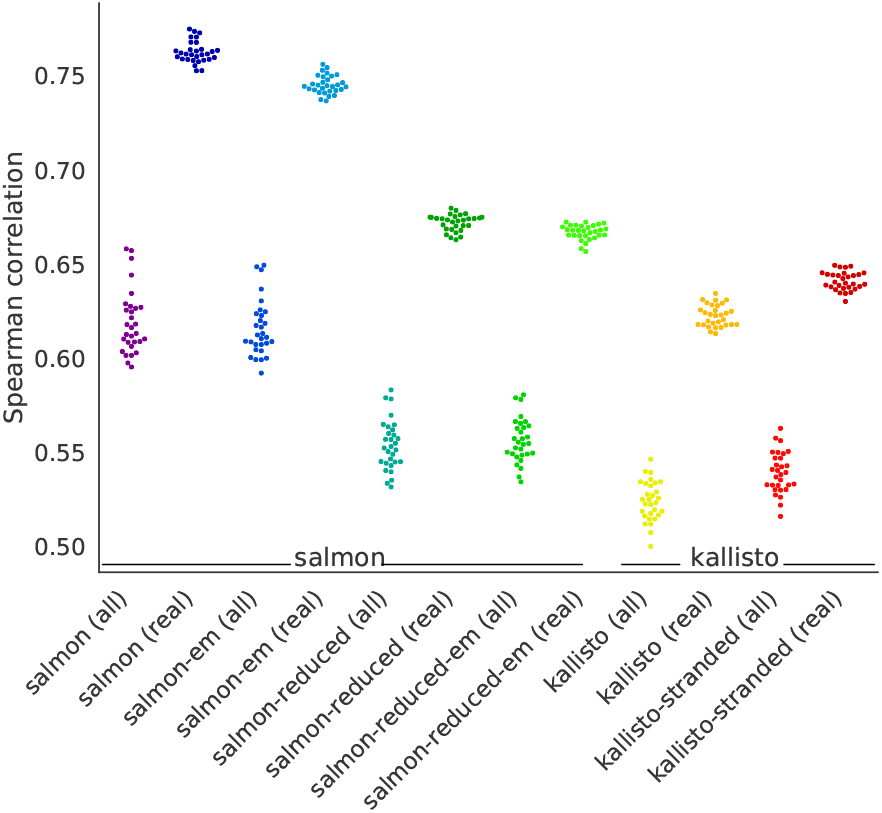
The distribution of Spearman ρ correlation coefficients for all method configurations on “all” (noisy) and “real” simulated samples.

**Fig. 2.**
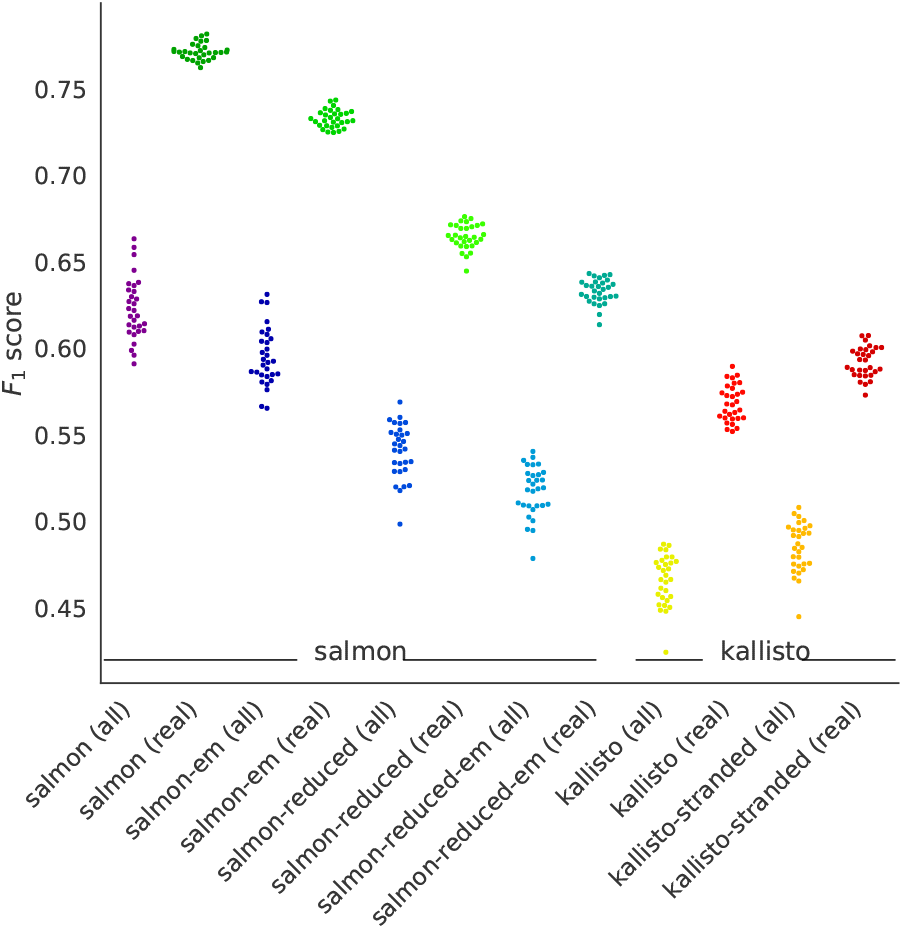
The distribution of F_1_ scores of all method configurations “all” (noisy) and “real” simulated samples.

**Fig. 3.**
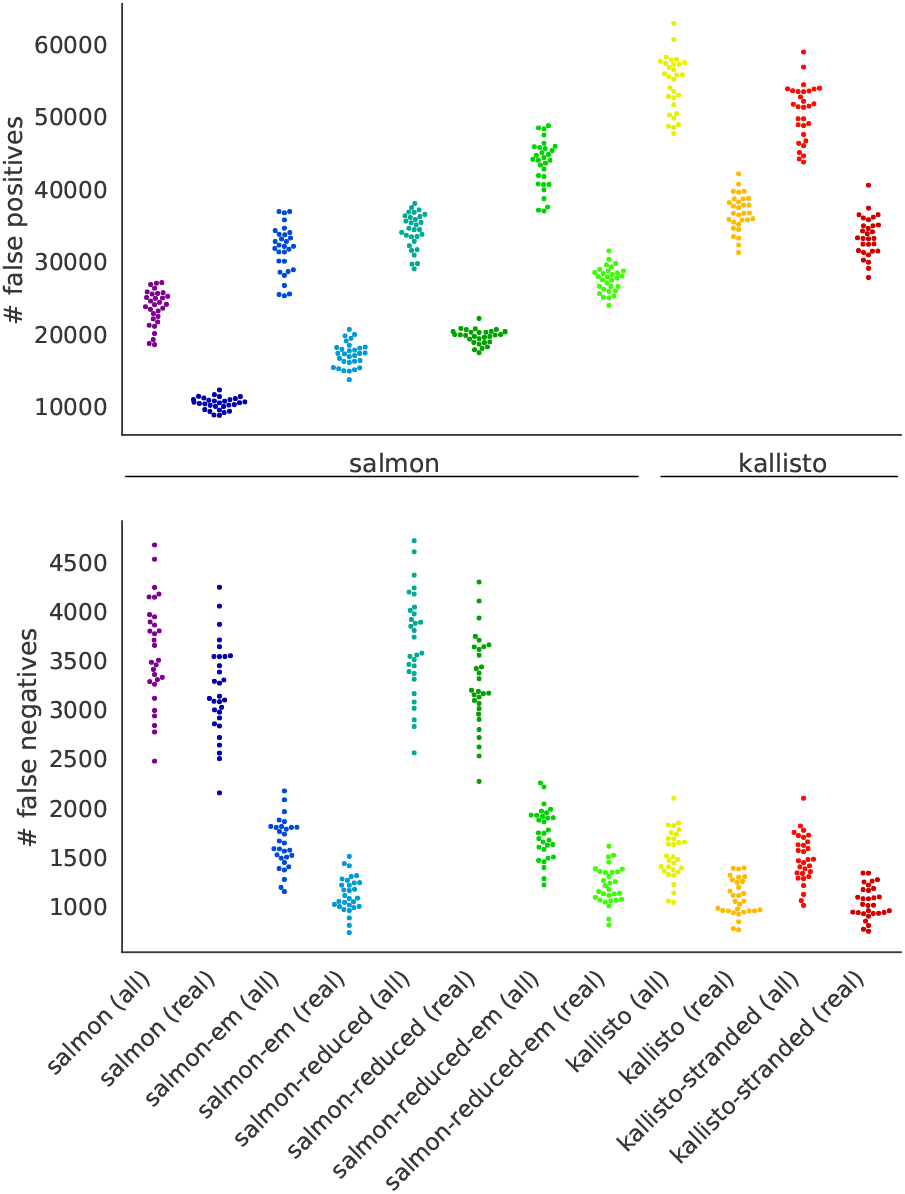
The distribution of false positive (top) and false negative (bottom) transcript counts of all method configurations on “all” (noisy) and “real” simulated samples.

**Fig. 4.**
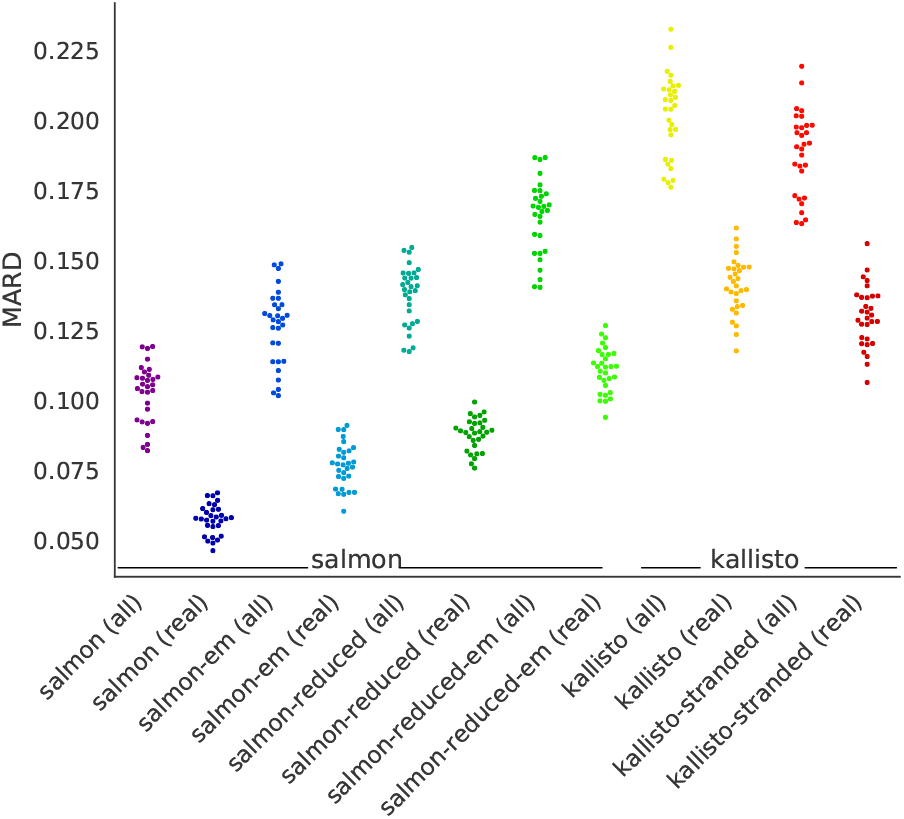
The distribution of mean absolute relative differences (MARDs) of all method configurations on “all” (noisy) and “real” simulated samples. A lower MARD indicates more accurate abundance estimates, with predicted fragment counts closer to true fragment counts.

The results presented in Figures 1 to 4 demonstrate that, regardless of the optimization algorithm used (VBEM, salmon’s default represented by the label “salmon”, or EM, represented by the label “salmon-em”), quantifying against a more comprehensive index, as suggested by Srivastava et al. (1), improves the accuracy of the estimation for both “real” and “all” samples in terms of Spearman correlation, F_1_ scores and mean absolute relative differences (MARDs). When operating with the reduced index, salmon provides somewhat more accurate predictions than kallisto on these simulated datasets, but both methods are considerably less accurate than the configuration of salmon that includes the expanded index. Also, as expected, compared to the corresponding “all” samples, all configurations provide more accurate estimations for samples from the “real” category, as the “real” samples include less transcriptional noise. The results also demonstrate that using the stranded option in kallisto improves the accuracy of it estimates in these samples under essentially all metrics, as is expected.

It is worth noting that neither the “real” nor “all” simulations represent pristine simulations of the type that are often performed to assess the internal consistency of quantification tools with their assumed generative model. That is, even though the “real” simulated samples do not include fragments arising from what is deemed as transcriptional noise, they will include fragments that arise from transcripts assembled in a tissue that match the intron chain of an annotated transcript but which may have alternative initiation or termination sites. This type of transcriptional diversity — ascertained from the experimental samples (8) — is designed to be representative of what is observed in experimental data, though it is often not fully represented in the annotation and can be highly tissue-specific.

When we examine the number of false positives (FPs) (annotated transcripts with estimated read counts > 0, but from which no fragments were simulated) and false negatives (FNs) (annotated transcripts from which fragments were simulated but with estimated read counts of 0) produced by each configuration (Fig. 3), we observe certain clear trends. While we evaluate FPs and FNs in terms of read count rather than TPM, and are using slightly newer versions of both salmon and kallisto, the trends we find among the methods and sample types assessed agree with the results reported by Varabyou et al. (2).

As has been discussed in previous literature (9), the variational Bayesian inference procedure, when used with a small prior, promotes sparse solutions. This can be seen in a simultaneous increase in the average number of FNs among configurations using the VBEM algorithm coupled with a considerable decrease in the average number of FPs. For example, when using the full index and assessing the results across the “all” samples, the VBEM configuration of salmon exhibits an average of ~ 3,582 FNs compared to the ~ 1,626 produced by the EM algorithm. However, the VBEM configuration exhibits an average of ~ 23,486 FPs compared to the ~ 31,470 produced by the EM algorithm. While the absolute number of FPs and FNs is lower across the “real” samples than “all” samples, the same trend persists. We observe that this trade-off between FPs and FNs can be adjusted by altering the prior applied to transcript abundances in the VBEM algorithm (data not shown).

When using the EM algorithm, across the “all” samples, all methods exhibit a similar number of false negatives (FNs), with kallisto-stranded exhibiting the fewest at an average of ~ 1,473 FNs and salmon-em exhibiting an average of ~ 1,626. However, kallisto exhibits the largest average number of false positives (FPs) at ~ 54,427, followed by kallisto-stranded at ~ 50,462. Meanwhile, among the EM variants of salmon, salmon-em exhibits ~ 31,470 FPs and salmon-reduced-em exhibits ~ 43,321. These data show that the EM inference procedure tends to reduce the number of FNs at the cost of increasing the number of detected FPs, while across methods, salmon maintains a lower false positive rate. Finally, as expected, regardless of the inference algorithm being used, the false positive rate is reduced when salmon’s full index is used rather than the reduced index.

## Conclusions

Depending on the transcriptional state and tissues in which they are measured, many RNA-seq samples will contain sequenced fragments arising from sequences not fully described by the annotated transcriptome, or from areas outside of the annotated transcriptome completely. As demonstrated by Varabyou et al. (2), this is true even in well-annotated organisms like human. Since annotation-based methods for transcript quantification rely explicitly upon a collection of annotated transcripts that are assumed to be complete, their accuracy will inevitably be affected by fragments that are incompatible with the annotated transcript models or that arise from other genomic loci, especially when those fragments can feasibly be mapped or aligned to annotated transcripts as well. Yet, the inherent difficulty of accurately assembling a per-sample transcriptome from short-read RNA-seq motivates the continued use of reference-based approaches when a “reasonably complete” transcriptome has been annotated or assembled for the samples being analyzed.

In this note, we have demonstrated that accounting for sequenced fragments originating from unexpected loci, as proposed by Srivastava et al. (1), improves the robustness and accuracy of reference-based transcript quantification compared to methods that consider only the annotated transcriptome. Using the simulation framework introduced by Varabyou et al. (2), we evaluate the quantification of transcript abundance in simulated RNA-seq samples when reads arise from mostly annotated loci (but with potentially tissue-specific transcription start and termination sites — *i.e*. the “real” simulated data), as well as when many different types of transcriptional noise (splicing, intronic and intergenic — *i.e*. the “all” simulated data) are present.

As expected, all tested methods perform best in the absence of transcriptional noise. However, considering a broader set of loci leads to improved accuracy under multiple metrics and under all types of simulated data. The improvements in accuracy can be substantial. Moving forward, it remains a challenge to continue to study the properties that arise in experimental RNA-seq samples that are not adequately modeled in simulation, as well as how quantification methods can best be improved to account for such properties. For example, while these simulations focus on transcriptional diversity and transcriptional noise, they do not account for certain biases, such as sequence specific and fragment-GC biases, known to be present in experimental RNA-seq data (10, 11). However, the results presented in (1, 2, 7) and in this manuscript suggest that one important aspect of accurate quantification will be properly determining when sequenced fragments, despite exhibiting sequence similarity to annotated transcripts, are most likely to have arisen from a transcribed sequence that differs from the annotation. In addition to the improvements introduced by Srivastava et al. (1), and in lieu of full transcript assembly, both transcript quantification over splicing graphs (12) and determination of optimal abundance ranges implied by non-identifiability in the underlying inference problem (13) are promising directions to pursue.

## Methods

We assessed two different annotation-based quantification tools (salmon (3) under 4 different configurations and kallisto (6) under 2 different configurations). Due to the fundamentally different nature of StringTie2 (4), which is capable of assembling and quantifying novel transcripts or transcript variants (with different initiation or termination sites), we focus only on the annotation-based quantification tools in this note. Further, we focus only on the more challenging problem of transcript-level quantification.

We evaluated salmon (v1.4.0) with the index prepared, with decoy sequence, as recommended by Srivastava et al. (1) under both the default inference settings (*i.e*. the variational Bayesian EM (VBEM) algorithm) and also using the traditional EM algorithm, providing salmon with the `--useEM` flag; these methods are denoted “salmon” and “salmon-em” respectively. The EM variant of salmon is included to aid in assessing how the specific inference method affects quantification accuracy, separately from the scope of the indexed reference. We also considered the same inference variants using a reduced index that consisted of only the annotated transcripts (as salmon was run in the Varabyou et al. (2) paper); these methods are denoted as “salmon-reduced” and “salmon-reduced-em”. In all cases, salmon removed 702 transcripts during indexing as exact sequence duplicates; if any such transcript is expressed in a sample, it will necessarily be considered as a false negative. Finally, we consider two different settings for the kallisto quantifier (v0.46.2), using the parameters provided in the Varabyou et al. (2) paper, and also explicitly providing the `--fr-stranded` flag (since these simulated samples are all stranded); these methods are denoted as “kallisto” and “kallisto-stranded”.

All methods were assessed using 60 separate simulated samples, corresponding to the 30 “real” samples and 30 “all” samples described in Varabyou et al. (2). The “real” samples consist of samples simulated across 3 different tissues (10 samples each), where the fragments are simulated from StringTie2 (4)-assembled transcripts in each tissue that matched — as determined by their intron chains — annotated transcripts from the CHESS 2.2 database (14), which was used as the reference annotation in all experiments. In addition to these simulated fragments, the “all” samples also include fragments simulated from different types of transcriptional noise (intronic, intergenic and splicing) detected in the experimental samples used to seed the simulations.

The reported metrics were computed as described below. False positives (FPs) are transcripts from which no simulated fragments were produced, but which were predicted to be expressed by a method. False negatives (FNs) are transcripts from which simulated fragments were produced, but which were predicted not to be expressed by a method. True positives (TPs) are transcripts from which simulated fragments were produced, and which were predicted to be expressed by a method. The F_1_ score is computed as (TP)/(TP +0.5(FP + FN)). The Spearman correlation coefficient was computed over the true and predicted fragment counts for all quantified transcripts using the implementation of pandas (15). Finally, the mean absolute relative difference (MARD) was computed over the true and predicted fragment counts for all M annotated transcripts using the following equation:

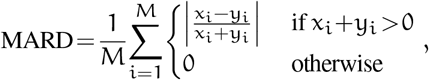

where x_i_ is the true number of fragments simulated from transcript i and y_i_ is the number of fragments estimated to arise from transcript i according to the quantifier being evaluated.

The scripts used to produce the results presented in this manuscript, based on the data of Varabyou et al. (2), are available at https://github.com/COMBINE-lab/quant-tx-diversity. The scripts make use of GNU Parallel (16). The repository also contains the Jupyter (17) notebook used to produce the plots, which makes use of seaborn (18), matplotlib (19), pandas (15), scipy (20) and numpy (21). The quantification results are deposited at https://doi.org/10.5281/zenodo.4437868.

